# The BDNF/TrkB pathway in Somatostatin-expressing neurons suppresses cocaine-seeking behaviour

**DOI:** 10.1101/2024.08.12.607442

**Authors:** Giuliano Didio, Teemu Aitta-aho, Juzoh Umemori, Eero Castrén

## Abstract

Cocaine addiction is a highly debilitating condition consisting of compulsive self-administration and seek for the substance of abuse, and its most challenging feature is the high rate of relapse. Addiction and relapse share similarities with neural plasticity which acts through the Brain-Derived Neurotrophic Factor and its receptor TrkB. Somatostatin (SST) expressing interneurons are involved in neuronal plasticity and are important in modulating cocaine-seeking behaviour in mice. We tested the role of TrkB in Somatostatin (SST)-expressing neurons in the extinction of cocaine-seeking behaviour, using mice in which TrkB has been knocked out specifically in SST neurons. We have observed that in these mice, once a cocaine-conditioned place preference is acquired, its extinction through seven days of extinction training is impaired, showing how this process relies on neural plasticity in SST neurons. When we promoted plasticity during extinction training using a light-activable TrkB in SST neurons in the prefrontal cortex of cocaine-conditioned mice, relapse of cocaine-seeking was prevented. Our data identify the critical role of TrkB-mediated plasticity within SST neurons in the extinction of and relapse to cocaine addiction.

## 1. Introduction

Addiction is a debilitating condition characterized by compulsive self-administration and a relentless pursuit of the substance of abuse and its most challenging aspect is the high rate of relapse, even after prolonged periods of abstinence (Brandon, Vidrine, and Litvin 2007). In 2010 alone, European Union countries collectively spent 7.6 billion euros on hospital treatments for addiction, which highlights the significant societal and economic impact of this condition (Lievens, Laenen, and Christiaens 2014). The neurobiological underpinnings of addiction involve several key brain areas, including the ventral tegmental area (VTA) during the binge/intoxication stage, the amygdala for the withdrawal/negative affect stage, and the orbitofrontal cortex, medial prefrontal cortex (mPFC), striatum, nucleus accumbens (NuAcc), hippocampus, and amygdala during the preoccupation/anticipation stage (Koob and Volkow 2010).

Similarities between the development of addiction and neural plasticity suggest a complex interplay between the brain’s adaptive mechanisms and the compulsive behaviors associated with addiction (Jones and Bonci 2005). The Brain-derived Neurotrophic Factor (BDNF) and its receptor neuronal receptor tyrosine kinase-2 (NTRK2, TrkB) are considered the main molecular switches of neural plasticity (Begni, Riva, and Cattaneo 2017), in fact deletion of BDNF in mice impairs LTP in hippocampal slices(Korte et al. 1995), and this effect can be rescued with long BDNF administration (Patterson et al. 1996). BDNF acts by binding to its receptor TrkB, activating it’s tyrosine kinase activity (Klein et al. 1989, 1991; Minichiello 2009) which in turn activates a cascade of secondary messengers, eventually leading to the phosphorylation of the transcription factor CREB, and promotion of plasticity-related genes (Finkbeiner et al. 1997; Minichiello et al. 2002). The BDNF/TrkB pathway is involved not only in LTP but also in neurogenesis, synaptic transmission and learning and memory (Bath, Akins, and Lee 2012; Bramham and Messaoudi 2005; Kang and Schuman 1995; Lee, Duan, and Mattson 2002; Li et al. 2008; Poo 2001).I

Increasing evidence is showing the involvement of BDNF/TrkB in the development and treatment of drug addiction (Jones and Bonci 2005; Nestler 2005).For example, it has been shown that the BDNF/TrkB pathway is necessary for the behavioral effects usually observed after cocaine administration (Hall et al. 2003). Mice with a heterozygous BDNF knockout displayed a lower cocaine-induced hyperlocomotion and showed a lower rate of cocaine-conditioning place preference (Hall et al. 2003). Moreover, in rats trained to self-administer cocaine and then exposed to long drug withdrawal, cocaine craving increased over time and BDNF (but not NGF) increased in VTA, NuAcc and amygdala as the time of drug withdrawal increased (Grimm et al. 2003). In rats after 4h of cocaine self-administration, there is an increase in the BDNF levels in NuAcc, and infusion of BDNF or BDNF-antibody in the NuAcc, increased or decreased respectively the cocaine self-administration and relapse (Graham et al. 2007). TrkB seems to be necessary for the development of cocaine-conditioned place preference (Crooks et al. 2010) and for the beneficial effects of environmental enrichment in preventing relapse of a pre-learned cocaine self-administration (Hastings et al. 2020). TrkB is widely expressed in the brain and in most of neurons, therefore the evidence mentioned so far does not clarify the role of single types of neurons in the regulation of addiction.

In the last years inhibitory neurons have gained attention as possible key players in addiction and other mental illnesses (Ostroumov and Dani 2018). SST-neurons are the second largest group of inhibitory neurons, representing 30% of the interneuronal population, present in all cortical layers (except for layer 1), they are generally classified as Martinotti and non-Martinotti cells (Tremblay, Lee, and Rudy 2016), with the majority being Martinotti-cells, characterised by dense dendritic trees and long translaminar axons (Riedemann 2019). SST-neurons target the distal dendrites of pyramidal cells and, interestingly, other inhibitory neurons like Vasoactive intestinal polypeptide (VIP)-expressing neurons and Parvalbumin (PV)-expressing neurons (Pfeffer et al. 2013; Riedemann 2019; Xiao et al. 2020), making these cells able to influence both excitation and inhibition in the network. Moreover SST-neurons seem to be involved in network plasticity, often through the inhibition of PV-neurons (Cummings and Clem 2020; Sadahiro et al. 2020; Tang et al. 2014).

SST-neurons in the Nucleus Accumbens have a bi-directional control over both the cocaine-induced hyperlocomotion and the cocaine-conditioned place preference (Ribeiro et al. 2018). Considering the strong involvement of neural plasticity and SST-neurons in the regulation of the addictive behaviour we decided to study the specific role of the BDNF/TrkB pathway in SST-neurons. With the use of mice expressing a heterozygous knockout of the *TrkB* gene (SST-TrkB-hCKO mice) and the conditioned place preference (CPP) paradigm (Itzhak and Martin 2002), we observed that while the *wt* group successfully reduced its cocaine-conditioning after a week of extinction training, the SST-TrkB-hCKO mice did not. However, both groups showed cocaine-induced reinstatement when tested a month later. We then specifically activated TrkB in SST-neurons in the Infralimbic cortex (ILCx) during the extinction training, by using, a sensitive type of optically activatable TrkB (E281A)(Hong and Heo 2020) with a mouse line expressing the *Cre* recombinase only in SST-neurons, and found that TrkB activation prevented relapse. This research provides evidence for the role of TrkB-mediated neural plasticity and SST-neurons in the extinction of a cocaine-conditioning and in the suppression of relapse. Our findings highlight the potential of targeting the BDNF/TrkB pathway in SST-neurons as a promising therapeutic strategy to consolidate extinction and reduce the risk of relapse in addiction treatments.

## 2. Materials and methods

### Generation of mice

Heterozygous TrkB knock out mice specifically in SST-neurons (SST-TrkB-hCKO) were generated by mating mice from a mouse line expressing the Cre recombinase in SST neurons (SST^Cre/Cre^; SST-IRES-Cre, RRID:IMSR_JAX:013044, Jackson Laboratory) with mice from a heterozygous line carrying floxed alleles of *Ntrk2*, the gene encoding for TrkB (TrkB^flx/wt^) (Minichiello et al. 1999) and *wt* littermate as controls. For the optogenetic experiment, we crossed SST^Cre/Cre^ with *wt* mice to generate SST^Cre/wt^ mice. The mice were kept in IVC cages with food and water *ad libitum*, with a 12-hours light/dark cycle, with light turning on at 6:00 am.

### Conditioned Place Preference (CPP)

The conditioned place preference (CPP) setup involved a Plexiglas box measuring 460×460×400mm and included removable floor panels of very different materials (smooth metal plates with holes (Type A) and rough plastic plates (Type B)). Each panel covered half of the box’s floor, requiring two panels to cover the entire surface. This allowed 4 possible combinations of floor configuration: AA; BB; AB; BA. The box was integrated into a system utilizing infrared signals to track the mouse’s position in real time, and a camera positioned above the box recorded the tests. To record the animals position we used the software TSE Multi Conditioning System.

The behavioural paradigm used for the CPP (**Fig.1**) test was:

**Fig. 1:**
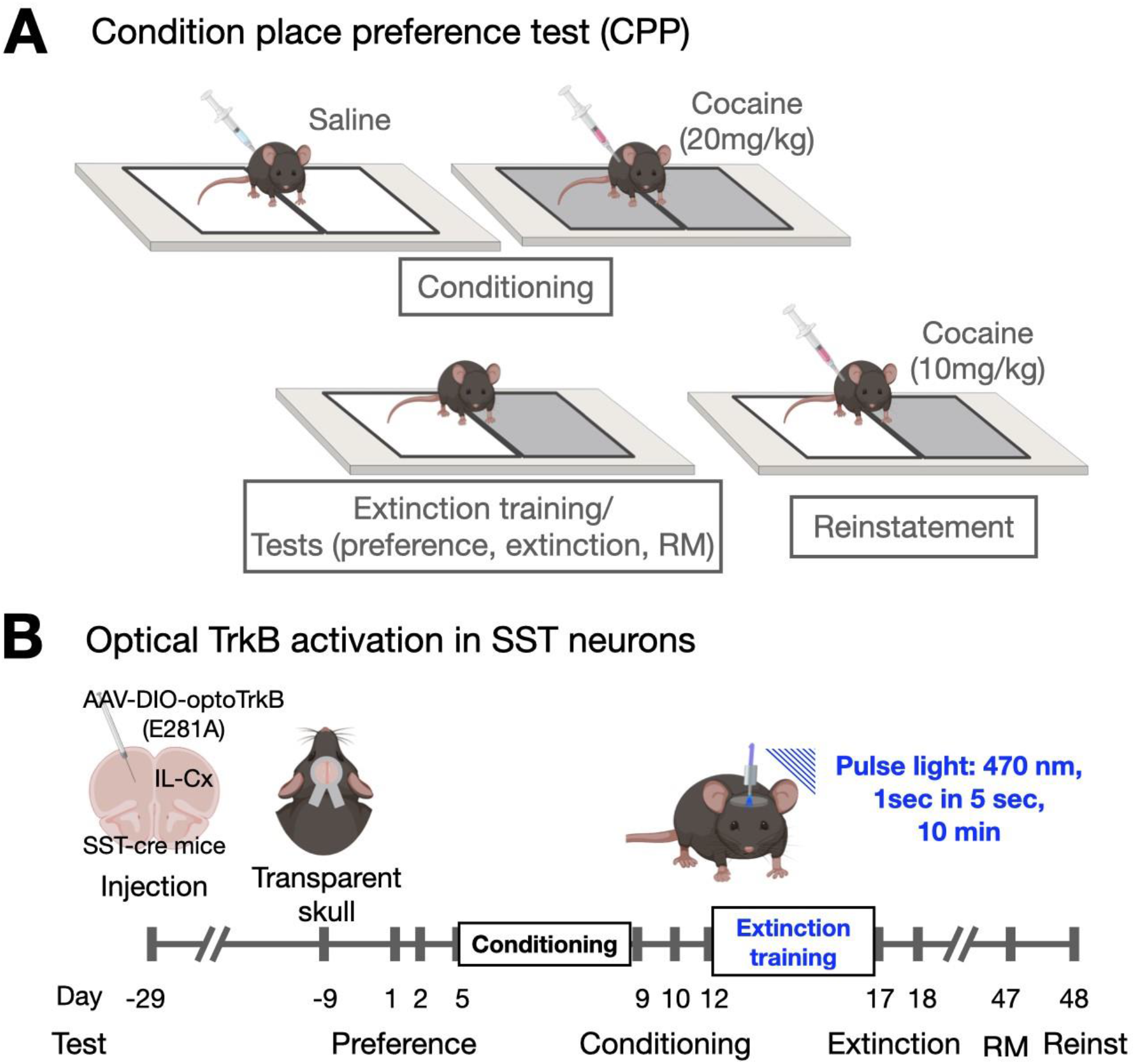
Schematic representation of the experimental design. **A**. Schematic representation of the CPP paradigm. Every mouse was exposed to a control conditioning involving saline injection and exposure to the “control floor pattern”, and then expose to the cocaine conditioning involving a cocaine injection and exposure to the “cocaine floor pattern”. When exposed to both floor patterns, a conditioned mouse would spend most the time on the cocaine floor pattern. **B**. Experimental design of the optogenetic activation of TrkB during CPP extinction. One month before the CPP, the mice were transduced with an AAV vector carrying optoTrkB. Three weeks later the mice underwent a procedure to make the skull transparent and to implant a metal holder above the IL-Cx. 9 days later the mice underwent the CPP paradigm. During each phase of the CPP, the mice had an optic fibre clipped magnetically to their metal holder. The light was turned ON only during the seven days of extinction training. Image created with Biorender.com

#### Day 1: Pre-test 1

Mice were injected with saline and placed in the box with Type A flooring on one half and Type B on the other for 20 minutes.

#### Day 2: Pre-test 2

Mice received another saline injection and were again exposed to Type A flooring on one side and Type B on the other for 20 minutes.

During these phases, we checked the mice’s initial preferences between the two floor types. Generally, preferences observed in Pre-Test 1 were consistent with those in Pre-Test 2. If results differed, we used the outcomes of Pre-Test 2. The results from these tests helped ensure that cocaine treatment was always paired with the less preferred floor type, while control treatment (saline injection) was associated with the preferred floor type.

#### Days 5-9: Conditioning

In the morning the mice underwent control treatment and were placed on the corresponding floor type. In the afternoon, they received cocaine treatment (20 mg/kg dissolved in saline, IP injection) and were placed on the corresponding floor type. The sessions lasted 15 min each.

#### Day 10: Conditioning Test

Mice were injected with saline and exposed to Type A flooring on one side and Type B on the other for 10 minutes.

This test assessed the level of conditioning. Mice were considered “conditioned” if they spent more than 50% of the time on the side associated with cocaine. Those that did not meet this criterion were excluded from the experiment.

#### Days 12-17: Extinction Phases

Mice received saline injections and were exposed to Type A flooring on one side and Type B on the other for 10 minutes.

Repeated exposure to both floor types in the absence of cocaine typically reduced preference until a roughly equal (50-50) preference was reached, indicating no preference.

#### Day 18: Extinction Test

Mice received saline and were exposed to Type A flooring on one side and Type B on the other for 10 minutes.

We used this test to determine the success of the extinction training.

#### Day 47: Remote Memory Test

Mice received a saline injection and were exposed to Type A flooring on one side and Type B on the other for 10 minutes.

We used this test to measure the long-term effects of extinction.

#### Day 48: Reinstatement Test

Mice were given a half dose of cocaine (10 mg/kg dissolved in saline, IP injection) and exposed to Type A flooring on one side and Type B on the other for 10 minutes.

This test models “relapse” (Shaham et al. 2003) and relies on the persistence of the neural association between the cocaine stimulus and the cocaine-associated floor type.

### Open Field Test

The Open Field box consisted of a 30 × 30 cm square arena provided by Med Associates, with transparent walls and a white smooth floor. Infrared sensors installed in the arena tracked both horizontal and vertical movements, recording the total distance travelled throughout the 10-minute test period. The arena was illuminated to approximately 150 lx. The peripheral zone was defined as a 6 cm wide strip along the edges of the walls.

### Immunohistochemistry

The brain slices were initially washed and permeabilized with a solution of PBS and 0.1% TritonX® (PBST) for 10 minutes at room temperature. Next, the samples were incubated in a blocking buffer composed of 3% Bovine Serum Albumin (BSA) in PBST for 1 hour at room temperature. Following the blocking step, I applied specific primary antibody solution (Rabbit anti-cFos, sc-052, SantaCruz (Dallas, Texas, USA), 1:300 in PBST) to the samples. The antibody was incubated with the samples overnight at 4°C. The next day, I washed the samples three times, 10 minutes each, with PBST at room temperature. Next, the samples were incubated with secondary antibody solution containing anti-Rabbit IgG antibody conjugated to Alexa647 fluorophore (Goat anti-rabbit IgG, Alexa647, Jackson Laboratory (Bar Harbor, Maine, USA), 1:800), diluted in PBST, for 2 hours at room temperature. After this step, the samples were washed three more times for 10 minutes each with PBST at room temperature, followed by a final wash with 0.1M phosphate buffer (PB 0) for 5 minutes. Finally, I mounted the samples on glass slides using Dako® mounting media and left them to dry overnight at room temperature, protected from light.

### Brain lysates analysis

The mice were euthanized with CO_2_, followed by cervical dislocation. We extracted the brain and isolated the mPFC while keeping everything on ice. The tissues were placed into 250 μL of lysis buffer containing NP40, vanadate as a phosphatase inhibitor, and a cocktail of protease inhibitors. We dissociated the tissue mechanically using a tissue disruptor. We measured the protein concentration using the BioRad DC Protein Assay, and the samples were then diluted with fresh lysis buffer to achieve uniform concentrations across all samples. Subsequently, the samples were denaturated using Laemmli buffer and placing them at 100 °C for 5 minutes. We run the samples onto NuPage™ 15-well gels (4-12% Bis-Tris gel) at 180 V until the 15 kDa marker exited the gel. The proteins were then transferred to a PVDF membrane using a blotting device at 100 V for 1.5 hours at 4°C. The membranes were washed with Tris Buffer Solution containing 0.001% Tween® 20 (TBST) for 5 minutes, then blocked with a blocking buffer (TBST + 3% BSA) for 1 hour at room temperature. Following blocking, the membranes were incubated with primary antibody solutions (p-CREB: #9198 Cell Signalling Technology, 1:1000; CREB: #4820 Cell Signalling Technology, 1:1000; HA: #2367Cell Signalling Technology, 1:500; all antibodies were diluted in TBST + 3% BSA) at 4°C for 24 to 48 hours. Next, the membranes were washed three times with TBST for 10 minutes each and incubated with secondary antibody solutions (HRP-conjugated secondary antibodies: Bio-RAD #1705045 or #1705047, diluted 1:10000 in TBST + 5% Non-Fat Dry Milk) for 2 hours at room temperature. After additional three washes with TBST for 10 minutes each, the membranes were treated with HRP substrate (Pierce™ ECL+, Thermo Scientific®) for 5 minutes at room temperature. The membranes were developed using a Syngene G:BOX device. The data was analysed using ImageJ.

### Image acquisition and analysis

The imaging of the slices was performed using a confocal microscope Zeiss LSM 780 or a widefield microscope Leica Thunder Imager 3D Cell Culture. The cFos analysis was performed by counting the positive nuclei and normalising the count on the area considered (ROI). The ROI was identified using the Allen Brain Atlas. The imaging was performed with the support of the Biomedicum Imaging Unit. We analyse the images using either Fiji ImageJ or Imaris 10.1.

### optoTrkB infection in the infralimbic cortex

We anesthetized the mice with Isoflurane and secured their heads in a stereotactic frame. We administered a subcutaneous injection of Carprofen (5 mg/kg) for pain relief, shaved the fur over the head of the mice, cleaned the area with Medetomidine, and made a single incision to expose the skull. The skull was cleaned with ethanol and air-dried. We drilled holes in the skull at the following coordinates: 1.78mm from Bregma, +/-0.7mm from the midline and -2.3 from the Dura Mater, so that with an injector carrying an inclination of 10° from the vertical axes, we could target specifically the infralimbic cortex. With a glass capillary loaded with the viral solution attached to an automatic injector (Stoelting Integrated Stereotactic Injector system) we injected the virus in the brain. The injection flow rate was set at 100 nl/min, and the capillary remained in place for 8 minutes following the injection. After the 8 minutes had elapsed, we carefully removed the capillary and closed the incision with histoglue (3M™ Vetbond™ Tissue Adhesive). The mice were placed on a heating pad to recover before being returned to their home cages.

### Transparent skull surgery and metal holder implant

This procedure was slightly adapted starting from a previous methods developed and published by our lab in 2017 (Steinzeig, Molotkov, and Castrén 2017). Two weeks after the viral infection, the mice were anesthetized with Isoflurane and secured on a stereotactic frame. After shaving the fur from their heads, we disinfected the exposed skin with 70% ethanol. The skin was then incised to reveal the skull, which was cleaned with acetone and sterile saline. To improve adhesion in the next step, I made a grid-like incision on the skull using a disinfected scalpel. I applied Loctite glue to the skull and allowed it to dry for roughly 5 minutes. Next, the metal holder (Steel430 ring with 8mm of outer diameter and 4 mm of inner diameter and 1mm of thickness. Custom design from Hubs.com) was attached with Loctite, ensuring the inner hole was positioned directly over the mPFC of the mouse. After another 5-minute drying period, we applied dental cement (Tetric EvoFlow) and solidified it using a UV lamp. Next, we prepared an acrylic mixture by combining colourless acrylic powder (EUBECOS, Germany) with methyl methacrylate liquid (Dentsply, Germany). I added a drop into the inner hole of the metal holder, then the mouse was placed on a heating pad to recover before being returned to its home cage.

### LED activation of optoTrkB

The setup involved optic fibers descending from above into the test box. We utilized a BioLED light source Control Module (Mightex) that was interfaced with the computer running the behavioral testing software, allowing for automated activation of the LED during the behavioural test. This module was connected to a single-color LED device (BLS Super High Power Fiber-Coupled LED Light Source, 470 nm Blue, Mightex). The LED device was linked to an optical connector wire, which connected to an optic fibre (custom ordered from ThorLabs) via a rotary joint (Rotary Joint Multimode Fiber Patchcord, 0.37 NA, 200μm Core, Mightex). These fibres terminated in a magnet, which clipped onto the metal holder positioned above the mouse’s PFC. The mice received pulses of 1s for the whole duration of the behavioural test (10 min) with an ITI of 5s.

### CPP with optoTrkB

We integrated the CPP paradigm with an optogenetic system. The mice equipped with a ferromagnetic-steel (steel430, company, and city?) metal holder around a transparent skull window for connecting to an optical fibre via a small, disk-shaped magnet that attached magnetically to the metal holder. This setup positioned the optical fibre above their prefrontal cortex (PFC). During the extinction phases, blue LED light was delivered through the optical fibre right over the PFC **(Fig.1)**.

### Statistical analysis

All the statistical tests were performed using GraphPad Prism. The data was tested using T-Test/Mann-Whitney, Wilcoxon test or a two-way ANOVA, depending on the number of variables considered and on the nature of the distribution of the data. Outliers were identified using the ROUT method (Motulsky and Brown 2006) with a Q value of 1, and any detected outliers resulted in the exclusion of that particular mouse from the specific analysis. A p-value of less than 0.05 was considered statistically significant.

## 3. Results

### TrkB in SST is necessary for the extinction of cocaine-induced CPP

For studying the role of neural plasticity in somatostatin expressing (SST) neurons in the development and extinction of cocaine-seeking behaviour, we performed the CPP paradigm using heterozygous conditional TrkB knockout mice (SST-TrkB hCKO) and wildtype littermate controls (*wt*) (**Fig.1A**). First, we conditioned the mice pairing cocaine and a distinguishable floor (plastic or metal). The mice showed a preference for the cocaine-associated floor and the proportion of conditioned mice seemed to be lower in SST-TrkB hCKO (43%) compared to *wt* littermate controls (70%), however, this effect did not reach statistical significance (X^2^-test, *wt* vs SST-TrkB-hCKO: p = 0.0579) (**Fig. 2B, Suppl. Fig. 1A-B**). To test the involvement of TrkB in SST neurons in the extinction of cocaine-seeking behaviour, the conditioned mice underwent an extinction training, with the aim of reducing the acquired preference by repeatedly exposing the mice to both the cocaine-associated and saline-associated floors without the drug. We observed that the *wt* control mice showed an expected reduction in preference (Paired T-test, p = 0.0028) (**Fig. 2C**). In contrast, the SST-TrkB hCKO mice continued to maintain their preference even after seven days of extinction training (Paired T-test, p = 0.6879). A month later, both groups underwent a remote memory test (RM) and a reinstatement test to assess relapse. Both groups demonstrated a preference for the floor pattern previously linked to cocaine, with SST-TrkB-hCKO even showing an increase in CPP (RM T-test, *wt* p = 0.0505; SST-TrkB-hCKO p = 0.022; Reinstatement T-test, *wt* p = 0.445; SST-TrkB-hCKO p = 0.263) (**Fig. 2 D, E**), indicating that the extinction did not occur in the SST-TrkB hCKO mice, and that in the *wt* mice, it was not lasting long enough to prevent relapse.

**Fig. 2:**
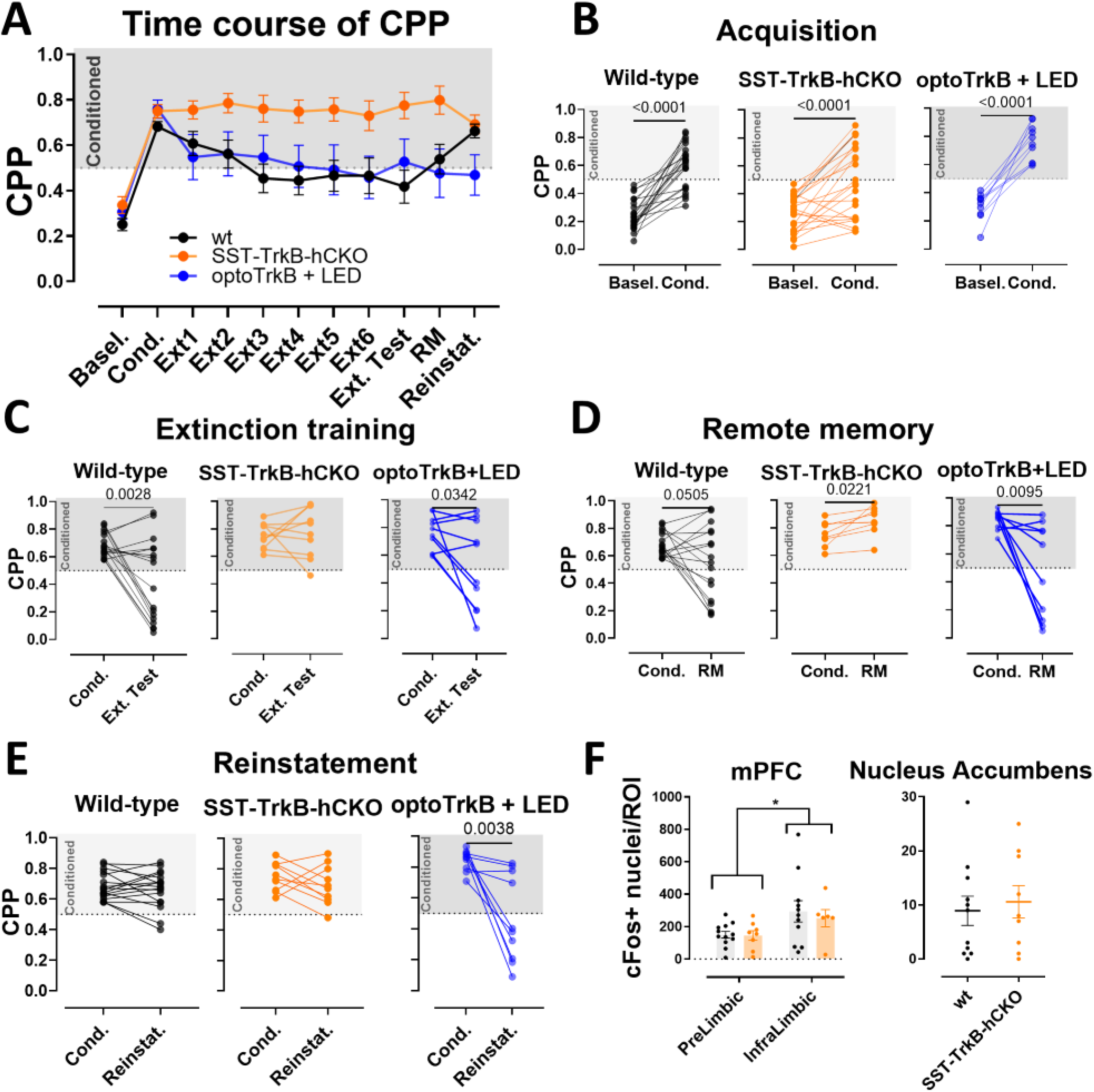
Results from CPP paradigm of *wt* (N=17, black), SST-TrkB-hCKO mice (N=10, orange) and mice infected with optoTrkB and exposed to LED light (N=10, blue). In figures B-F each dot represents a mouse. **A**. Time course of the whole experiment. **B**. Paired T-tests of the different groups between their baseline behaviour and their preference after 5 days of conditioning. Paired T-tests: *wt* p < 0.0001; SST-TrkB-hCKO p < 0.0001; optoTrkB + LED p < 0.0001. **C**. Preference between the post-conditioning test and after a week of extinction training. Paired T-tests: *wt* p = 0.0028; SST-TrkB-hCKO p = 0.6879; optoTrkB + LED p = 0.0342. **D**. Preference between the post-conditioning test and the remote memory test. Paired T-tests: *wt* p = 0.0505; SST-TrkB-hCKO p = 0.0221; optoTrkB + LED p = 0.0095. **E**. Preference between post-conditioning test and reinstatement test. Paired T-test: *wt* p = 0.445; SST-TrkB-hCKO p = 0.263; optoTrkB + LED p = 0.0038. **F**. Count of cFos-positive nuclei normalised on the area of ROI. Two-way ANOVA of mPFC: Genotype F (1, 33) = 0.229, p = 0.635; brain area F I (1, 33) = 6.650, p = 0.0146. Nucleus Accumbens T-test p = 0.688.

Since we observed a difference in the levels of extinction between the SST-TrkB-hCKO and the *wt* mice, we thought of checking whether there was any difference in the activation of neural networks associated with the promotion of extinction and inhibition of relapse, like the IL-Cx and the NuAcc (Bahi, Boyer, and Dreyer 2008; Chauvet et al. 2009; Hastings et al. 2020; LaLumiere, Niehoff, and Kalivas 2010; Nestler 2005; Peters, LaLumiere, and Kalivas 2008). One hour after the relapse test, we euthanized the mice and measured the levels of cFos, an immediate early gene that is expressed in response to recent neural activity (Hudson 2018) and found no significant differences in recent activation between the SST-TrkB hCKO mice and their *wt* littermates (**Fig.2 F**) both in the mPFC (2-way ANOVA mPFC, Genotype F(1, 33) = 0.229, p = 0.635) and in the NuAcc (T-test, p = 0.688). However, the 2-way ANOVA test in the mPFC highlighted a difference in the levels of cFos between the two brain areas forming the mPFC, suggesting a higher level of neural activation in the infralimbic cortex (IL-Cx) compared to the Prelimbic cortex (PL-Cx) (2-way ANOVA mPFC, brain area F(1, 33) = 6.650, p = 0.0146). This led us to hypothesise that neural plasticity, in SST neurons, in this very area could be manipulated to consolidate the extinction training network and prevent relapse.

### TrkB activity in SST neurons is sufficient to inhibit the reinstatement of cocaine-induced CPP

We hypothesized that enhanced neural plasticity in SST-neurons during extinction training might strengthen the extinction and prevent the relapse rate of cocaine-seeking behaviour. To regulate neural plasticity specifically in the SST neurons, we transduced a sensitive version of optoTrkB (optoTrkB (E821A) (Hong and Heo 2020; Lilja et al. 2022) (**Fig. 3A**) into the IL-Cx of SST-cre mice. Immunohistochemistry for the HA-tagged optoTrkB confirmed the correct location of optoTrkB and its colocalization with SST (**Fig. 3B, D**). LED exposure significantly increased pCREB levels in the IL-Cx lysates of these mice compared to unstimulated control mice (Mann Whitney test, p = 0.0043) (**Fig. 3C**), confirming the activation of the downstream pathway of BDNF/TrkB (Finkbeiner et al. 1997; Minichiello et al. 2002). We also observed a significant difference in the levels of pCREB between the unstimulated infected mice and uninfected controls (**Supp. Fig. 2E**), which may be produced by some spontaneous activation of TrkB upon overexpression (Koponen, Võikar, et al. 2004; Koponen, Lakso, and Castrén 2004; Schecterson and Bothwell 2010). We conducted a CPP experiment with SST-optoTrkB mice, and during the extinction training we exposed blue light (λ=470 nm, 1s exposure with 5s ITI, for 10 min) to activate optoTrkB in SST neurons (**Fig. 1A**). As expected, SST-optoTrkB mice exposed to LED developed a clear conditioning place preference as the *wt* mice did (Paired T-test, p < 0.0001) (**Fig. 2B**). The SST-optoTrkB showed a significantly decreased preference after six days of extinction training (Paired T-test, p = 0.0342) (**Fig. 2C**). When tested for remote memory, the mice seemed to be still benefiting from the effects of the extinction training (Paired T-test, p = 0.0095), showing low levels of cocaine-place preference (**Fig. 2D**). Most interestingly, the LED group showed no increase in freezing, in the reinstatement test for relapse (Paired T-test, p = 0.0038) (**Fig. 2 A; E**), suggesting an enhanced effect of extinction training with the optoTrkB activation. The unstimulated control mice also showed similar results as the stimulated mice in the extinction, remote memory, and reinstatement tests (Extinction Paired T-test, p = 0.0109; RM Wilcoxon, p = 0.1289; Reinstatement Paired T-test, p = 0.0128) (**Supp. Fig.2B-D**), which is consistent with the increase in pCREB baseline levels in the absence of LED stimulation and suggests that the spontaneous TrkB activation due to overexpression of TrkB in SST neurons in the IL-Cx might be sufficient to inhibit relapse. Notably, when comparing each mouse’s preference in the RM test to its own preference right after conditioning, only the LED group had a statistically significant shift in preference (LED-group, Paired T-test, p = 0.0095; no-LED group, Wilcoxon, p = 0.1289) **(Fig. 1D; Supp. Fig. 2C**), suggesting a more solid effect in the optoTrkB activated group.

**Fig. 3:**
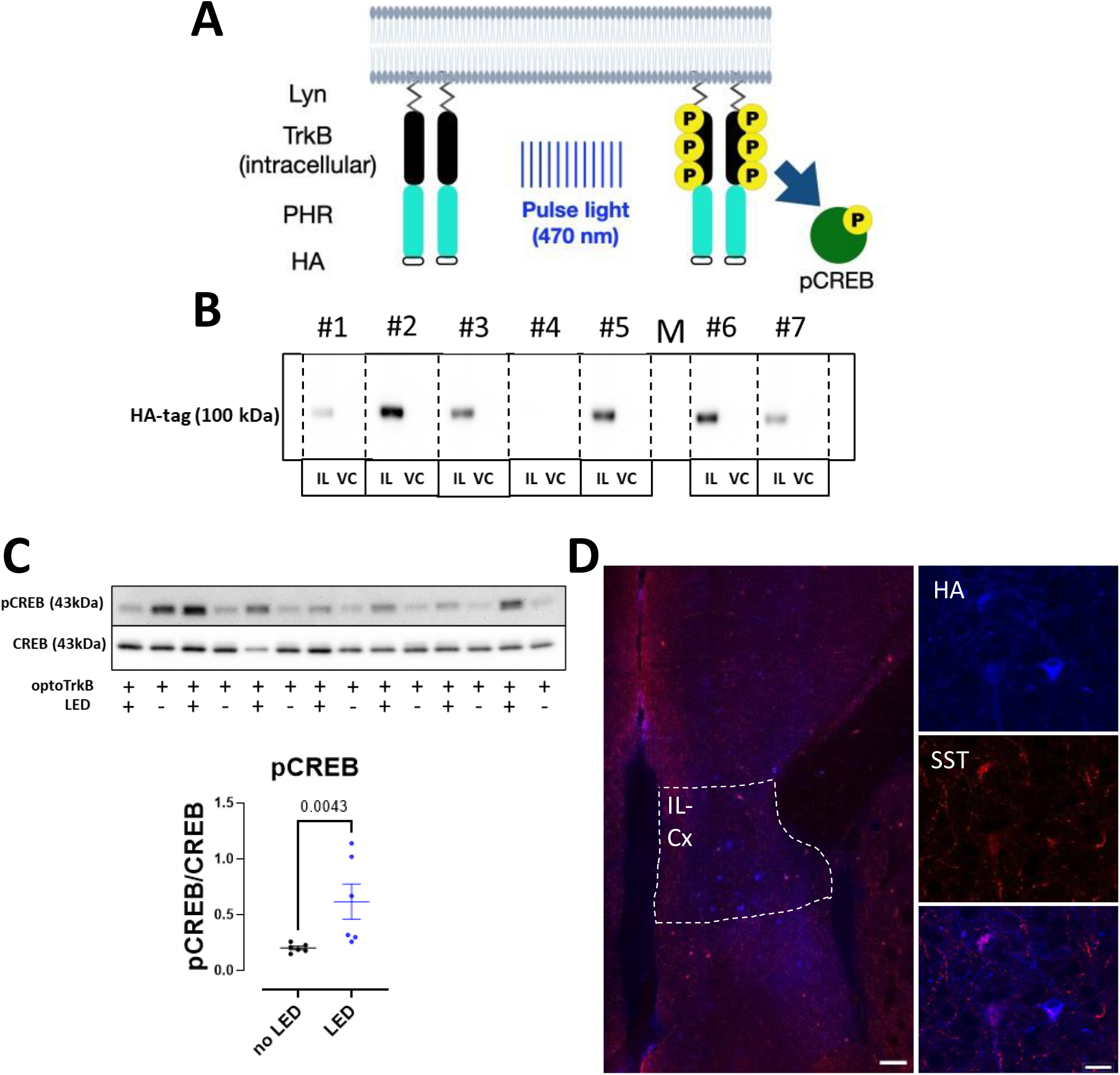
Assessment of optoTrkB E281A. **A**. Schematic representation of the structure of optoTrkB. The construct presents the intracellular domain of TrkB bound to a Photolyse Homology Region (PHR) and a HA-tag. The localisation of optoTrkB to the membrane is ensured by a Lyn sequence on its N-terminus. Upon exposure to blue light, the PHR domain promotes the homodimerization of the optoTrkB monomers, triggering the secondary messengers signalling cascade, leading to the phosphorylation of CREB **B**. Western blot with antibody anti-HA tag. Vertical dotted bands identify a mouse. For each mouse the infected area (IL-Cx) and a control area (Visual cortex, VC) is shown. The HA-positive band is present only in the infected brain area of each mouse. **C**. Western blot bands for phosphor-CREB and CREB and quantification. Mann Whitney test, p = 0.0043. **D**. Confocal images of infralimbic cortex of mice infected with optoTrkB E281A. the HA tag is shown in blue, SST is shown in red. The size bar in the low magnification image is 100 μm (on the left), while in the image with higher magnification (on the right) is 20 μm.

## 4. Discussion

In this study, we assessed the role of TrkB in SST neurons in the development and extinction of conditioning to cocaine. Our first approach consisted of impairing the BDNF/TrkB pathway in SST neurons of mice that underwent a CPP paradigm. The main observation was the complete lack of effectiveness of the extinction training on the SST-TrkB-hCKO mice that acquired the CPP. This suggests how TrkB in SST neurons is necessary for extinguishing such conditioning. However, the *wt* clearly benefited from the extinction training, showed a slight increase in preference one month after the extinction training and showed a full relapse when tested for reinstatement. This implies that the benefits of the extinction trainings were not long-lasting and did not delete the memory associated to the conditioning to cocaine. We observed no difference in the levels of cFos between the two groups one hour after the reinstatement test, neither in the IL-Cx nor in the NuAcc. However, when including in the analysis the comparison between cFos in the IL-Cx and in the Pr-Cx (brain area that promote cocaine-seeking behaviour (McFarland, Lapish, and Kalivas 2003), we saw a significantly higher amount of cFos nuclei/ROI in the IL-Cx. A higher activity in the brain area associated with extinction one hour after a test where the mice showed full relapse, might reflect a persistence of an “extinction” neural network that competes with the “cocaine-seeking” network. These observations made us hypothesise that if we were to promote plasticity in this “extinction” network, we would be able to prevent relapse.

To enhance neural plasticity in SST neurons within the IL-Cx, we used an optogenetic TrkB receptor. After assessing the correct location and activation of optoTrkB, we used it to consolidate neural networks created during the extinction training and prevent relapse. The observations that mice with optoTrkB activation during extinction training did not relapse suggests that TrkB activation in SST neurons during extinction training is sufficient to consolidate the extinction network and prevent relapse. Enhancement of behavioural flexibility has been obtained via different mean by researchers. For example it has been shown how long-term ketamine administration can promote fear erasure (Ju et al. 2017), injection of TrkB agonist in the IL-Cx promotes the extinction of CPP (Otis, Fitzgerald, and Mueller 2014), and the psychedelic psilocybin has been shown to promote behavioural flexibility in rats (Torrado Pacheco et al. 2023). Our lab has showed how manipulating TrkB specifically in inhibitory neurons can lead to increased neural plasticity and induce behavioural and network shifts that would otherwise happen only temporarily or wouldn’t happen at all. For example, in 2011 and in 2023 our group showed how the plasticity-inducing antidepressant drug fluoxetine consolidates the effects of a conditioned-fear extinction training, preventing fear renewal and recovery, and that this was dependent on TrkB in the amygdala and on TrkB specifically on parvalbumin (PV)-expressing inhibitory neurons (Jetsonen et al. 2023; Karpova et al. 2011). Also in 2021 we showed how activating optogenetically TrkB in PV neurons in the visual cortex of adult mice, it is possible to induce a shift of ocular dominance that usually happens only in young mice (Winkel et al. 2021). We called this induced juvenile-like plasticity “iPlasticity” (Umemori et al. 2018). Moreover, other research has further confirmed how manipulating TrkB and/or inhibitory neurons we can induce a heightened state of neural plasticity (Harauzov et al. 2010; Lensjø et al. 2017; Sale et al. 2007)

In this work we might be seeing a similar phenomenon. *Wt* mice already benefited from a CPP extinction training, without the need of boosting its BDNF/TrkB pathway, but this beneficial effect was not permanent, and relapse kicked in when the “cocaine-seeking network” was primed with a drug cue, suggesting the existence of two neural networks (cocaine-seeking network and extinction network) that would compete with each other. By boosting TrkB in the neurons relevant for the extinction of the CPP (the SST neurons), in the relevant brain area (the IL-Cx), during the relevant behavioural training (the extinction training), we selectively consolidated the “extinction network”, forcing the competition in favour of the extinction, thus inhibiting relapse.

Interestingly, even mice expressing optoTrkB E821A without LED stimulation showed reduced relapse. Given that both groups demonstrated strong cocaine-conditioned place preference after the conditioning, and that cocaine-induced reinstatement is a well-established and widely observed phenomenon (Aguilar, Rodríguez-Arias, and Miñarro 2009), our finding suggests that the expression of optoTrkB E821A alone may enhance neural plasticity. This hypothesis is supported by the observation that unstimulated optoTrkB-expressing mice had higher levels of pCREB than uninfected controls. Trk receptors are known to auto-dimerize when overexpressed (Schecterson and Bothwell 2010), and TrkB overexpression has been linked to increased neural plasticity *in vivo* (Koponen, Võikar, et al. 2004; Koponen, Lakso, et al. 2004). Therefore, our data suggests that heightened TrkB activity, through either optogenetic means or overexpression in SST neurons in the IL-Cx, coupled with a CPP extinction training, can support the extinction neural network, and prevent relapse.

## 5. Conclusion

In this study, we demonstrated that TrkB activity in SST neurons is essential for a successful extinction of cocaine-seeking behaviour in mice. Moreover, we found that by enhancing the BDNF/TrkB signalling pathway in SST neurons during extinction training promotes extinction and prevent relapse. As relapse is a major problem in addiction, these findings highlight the importance of neural plasticity, particularly in inhibitory neurons, in the treatment of addictive behaviours. Future studies are needed to assess the clinical implications of these observations to develop more effective interventions that focus on preventing relapse.

## 6. Acknowledgments

We would like to thank Sulo Kolehmainen for the research support and for genotyping the mice, Vootele Voikar, Nelli Koivisto and Aurora Hämäläinen for the constant support with the behavioural setups and the caretakers of the animal facility of the University of Helsinki. We thank Dr. Anni-Maija Linden for providing us the SST-cre mice used to generate our mice lines, and Dr. Jongryul Hong and Dr. Won Do Heo for providing the plasmid of the optoTrkB E281A. The behavioural experiments were carried out with the support of the Mouse Behavioural Phenotyping Facility supported by Biocenter Finland and HiLIFE. The presented research was supported by the Academy of Finland grants #2327192307416 and 347358, the Sigrid Jusélius foundation, Jane & Aatos Erkko Foundation, the HiLife Fellows program, Bilateral exchange program between the Academy of Finland and JSPS (Japan Society for the Promotion of Science).

## Ethical statement

The experiments were carried out in accordance with the European Communities Council Directive 86/6609/EEC and the guidelines of the Society for Neuroscience and were approved by the County Administrative Board of Southern Finland (License number: ESAVI/40845/2022).

## Author contributions

G.D., TA, EC and J.U. designed the experiment, G.D. carried out the experiments and the analysis. G.D., J.U. and E.C. wrote the manuscript.

---

**Suppl. Fig. 1:**
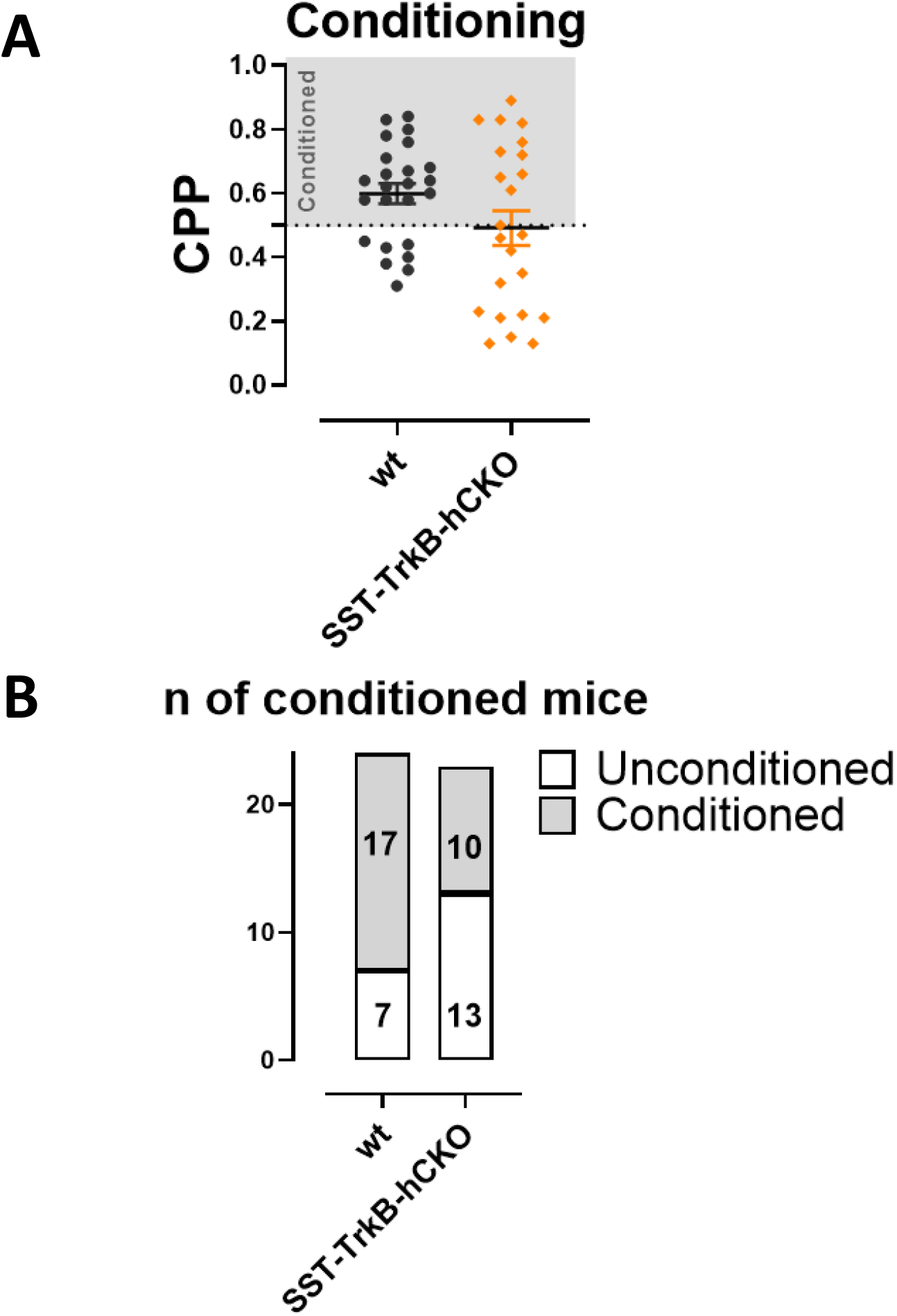
Data from conditioning test of the CPP between *wt* mice and the SST-TrkB-hCKO mice. **A**. Preference after five days of conditioning. T-test p = 0.0896. **B**. Number of mice that showed a preference for the cocaine-associated side above 0.5 (conditioned) and mice that showed a preference below or equal to 0.5 (unconditioned), between the two groups. X^2^-test, *wt* vs SST-TrkB-hCKO: p = 0.0579.

**Suppl. Fig. 2:**
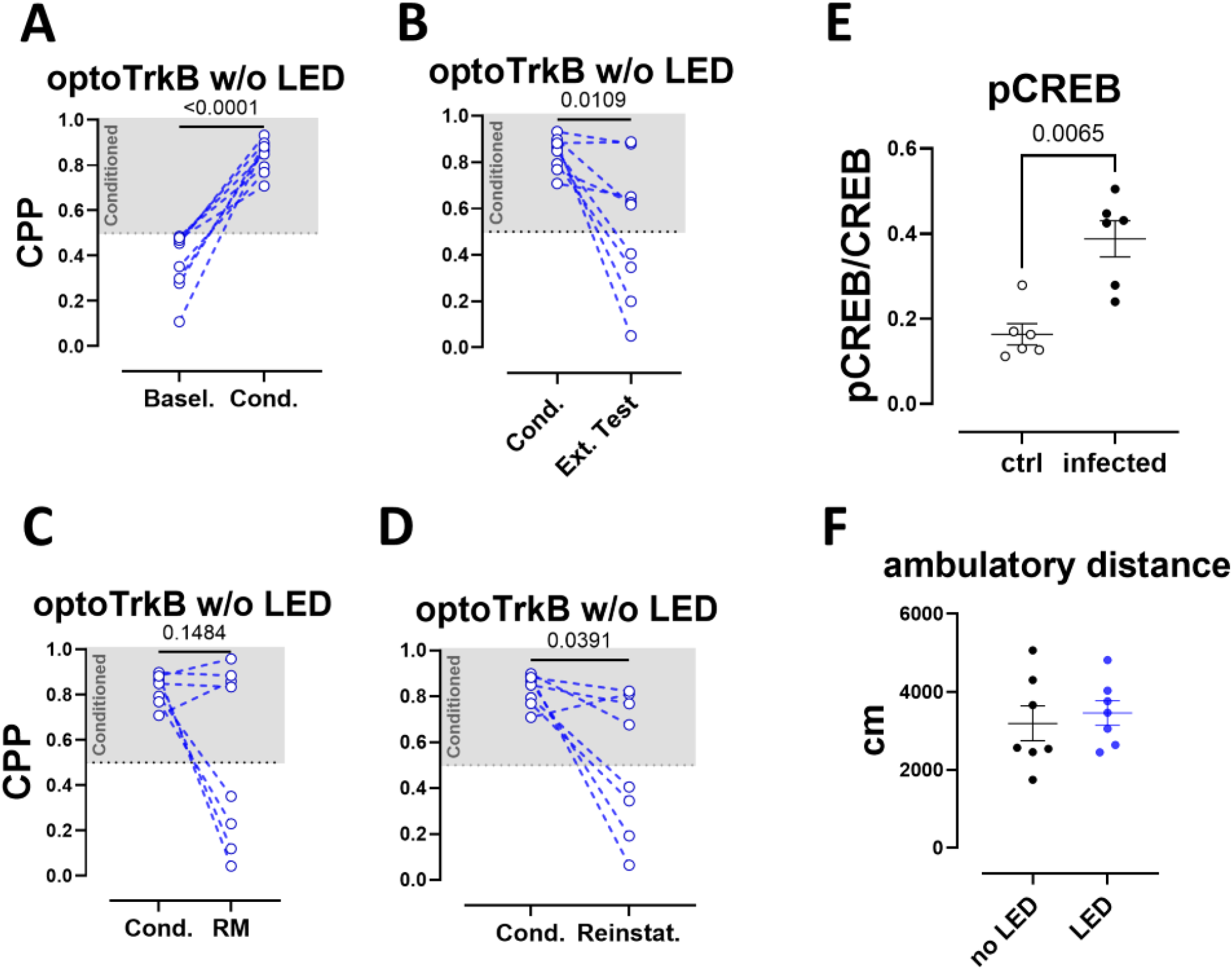
Preference rate of optoTrkB mice without LED stimulation (N=9). **A**. Preference before and after five days of conditioning. Paired T-test p < 0.0001. **B**. Preference rate between post-conditioning test and after seven days of extinction training. Paired T-test, p = 0.0109. **C**. Preference rate between post-conditioning test and remote memory test. Wilcoxon, p = 0.1289. **D**. Preference between post-conditioning test and reinstatement. Paired T-test, p = 0.0128. **E**. Analysis of western blot bands of pCREB/CREB between optoTrkB w/o LED mice and uninfected controls. Mann-Whitney test, p = 0.0065. **F**. LED activation of optoTrkB has no effect on baseline locomotion. Mann Whitney test, p = 0.6200.

